# A role for the anterior hippocampus in autobiographical memory construction regardless of temporal distance

**DOI:** 10.1101/2022.05.01.490212

**Authors:** Sam Audrain, Adrian W. Gilmore, Jenna M. Wilson, Daniel L. Schacter, Alex Martin

## Abstract

Mounting evidence suggests distinct functional contributions of the anterior and posterior hippocampus to autobiographical memory retrieval, but how these subregions function under different retrieval demands as memories age is not yet understood. Specifically, autobiographical memory retrieval is not a homogenous process, rather, it is thought to consist of multiple stages: an early stage of memory construction and a later stage of detailed elaboration, which may differently engage the hippocampus over time. In the present study, we analyzed data from 40 participants who constructed and overtly elaborated upon recent and remote memories in response to picture cues in the fMRI scanner. We previously reported a temporal gradient in the posterior hippocampus during the elaboration period of autobiographical retrieval, with posterior hippocampal activation observed for recent but not remote timepoints. Here, we consider the previously unanalyzed construction stage of retrieval, where participants searched for and selected a memory. We found no evidence of a temporal gradient during memory construction, instead observing strong anterior hippocampus activity regardless of memory remoteness. Our findings suggest a unique contribution of the anterior hippocampus to the construction process of autobiographical retrieval over time. These findings highlight that retrieval processes, which have yet to be considered in current models of systems consolidation, offer novel insights to hippocampal subregion function over time.

## Introduction

The human ability to mentally travel back in time to re-experience past events— “autobiographical” or “episodic” memory—is central to human cognition. This capacity helps give rise to our sense of self (Conway and Pleydell-Pearce, 2000), forms the basis for interactions around the dinner table (Mahr and Csibra, 2018), and likely supports our ability to envision future events (Tulving, 1985; Schacter and Addis, 2007). The hippocampus and other medial temporal lobe structures, through interactions with one another and the neocortex, are thought to be critical for both the formation and retrieval of autobiographical memories (Squire and Zola-Morgan, 1991). Indeed, among the most common questions still asked by memory researchers today is how the hippocampus supports memory retrieval.

However, converging lines of research suggest that investigating “the” role of “the” hippocampus may frame the question too broadly. Perhaps most notably, the characteristics of retrieved memories appear important in determining hippocampal contributions to recall. Results spanning decades have highlighted differing roles for the hippocampus in retrieving recent and remote memories, even though the basis for such change over time remains debated. One prominent model assumes a time-limited role for the hippocampus, with memories being “consolidated” to the neocortex over time (the “Standard Model of Consolidation”; Squire et al., 2015). Others have argued that the propensity of memories to degrade over time drives reductions in activity for remote memories, and that the hippocampus is always necessary for recalling richly-detailed memories (“Trace Transformation Theory”; Nadel and Moscovitch, 1997; Sekeres et al., 2018).

Joining the idea of memory properties themselves altering hippocampal responses are two additional lines of research. One describes heterogeneity along the long-axis of the hippocampus. Anterior and posterior regions possess distinct structural connectivity, functional connectivity, and task activation profiles (Poppenk et al., 2013; Barnett et al., 2019; Persichetti et al., 2021; Zheng et al., 2021). Evidence suggests that the posterior hippocampus supports the retrieval of fine-grained details whereas the anterior hippocampus supports coarser, gist-like memory features (Brunec et al., 2018; Sekeres et al., 2018; Audrain and McAndrews, 2020). In addition, there may be non-stationarity in the processes involved in retrieving autobiographical memories. Conway’s (2005) Self-Memory System framework proposes that autobiographical memory retrieval can be separated into two conceptually and temporally discrete stages. An early “construction” period involves the search and recovery of general aspects of a memory, whereas a later “elaboration” period involves a sustained recollection period wherein the distinct elements of an episode are re-experienced in detail (Conway, 2005).

Hippocampal contributions to autobiographical retrieval may therefore depend on several distinct factors. Studies with this in mind remain relatively rare yet have provided intriguing results. For instance, a recent fMRI investigation of effective connectivity in anterior and posterior hippocampal subregions during construction and elaboration stages found changes in the direction of information flow within and beyond the anterior and posterior hippocampus across retrieval stages (McCormick et al., 2015, 2018). However, simple univariate differences in hippocampal engagement were not observed. Similarly, Addis et al., 2007 associated anterior hippocampal activity with construction-related activity across several comparisons, but critically, not in their direct contrast of construction and elaboration stage activity. Neither study examined the effect of event recency on hippocampal engagement, although a subsequent re-analysis of the elaboration phase of the Addis et. al (2007) data indicated a temporal gradient for future events (distant future>near future) in the hippocampus (Addis and Schacter, 2008). In some of our own recent work, the roles of the anterior and posterior hippocampus during the elaboration phase of retrieval for recent and remote past events was investigated using overt in-scanner recall (Gilmore et al., 2021b). These results identified temporally graded activity in the posterior hippocampus, without consistent engagement of the anterior hippocampus. However, our analyses were restricted to the elaboration phase. Thus, these data provided no insights into how the construction phase might be supported by anterior or posterior aspects of the hippocampus or how it might be impacted by event recency. With this shortcoming in mind, we analyzed data from our previously published experiment (Gilmore et al., 2021b), but now focusing on activity during the memory construction stage of each trial (**Figure 1**).

**Figure 1.**
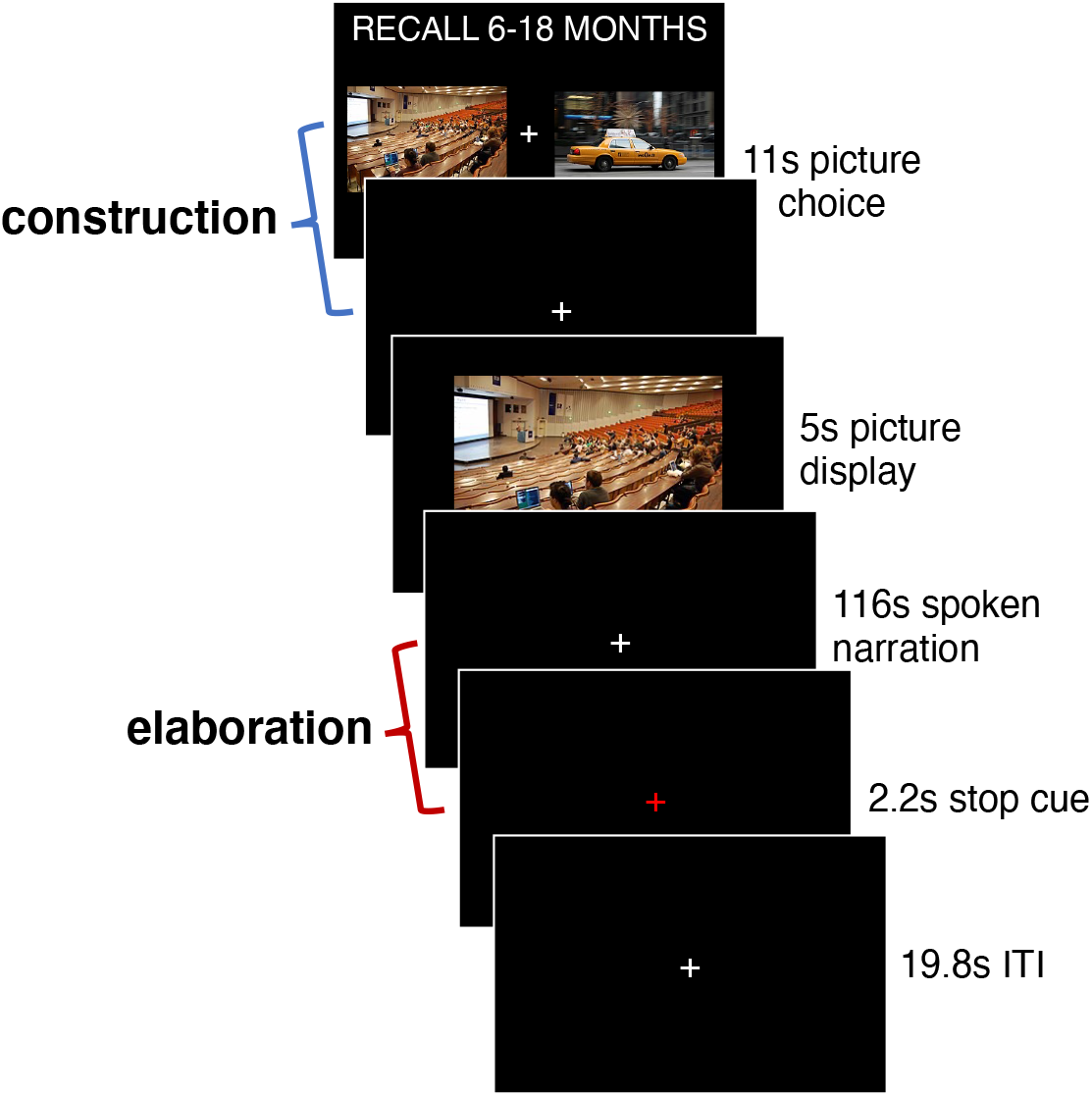
Trial structure of the autobiographical retrieval task. During the construction stage, participants selected their preferred picture cue to use to retrieve autobiographical memories from earlier in the day, 6-18 months ago, or 5-10 years ago. During elaboration, participants overtly described the retrieved memory in as much detail as possible. The trial structure for the control task was the same, except participants chose a picture (construction stage) to describe in detail (elaboration stage), without retrieving a long-term memory.

## Materials and Methods

### Participants

Data were collected from forty right-handed young adult participants (23 female; mean age = 24.2 years old). All were native English speakers with normal or corrected-to-normal vision and reported no history of psychiatric or neurologic illness. Informed consent was obtained from all participants and the experiment was approved by the NIH Institutional Review Board (clinical trials number NCT00001360). Participants received monetary compensation for their participation. For additional participant information, see Gilmore et al. (2021b).

### Experimental Design

#### Stimuli

Stimuli consisted of 48 images depicting complex scenes (i.e., people engaging in various activities in a specific location). Images were sized at 525 × 395 pixels (screen resolution: 1920 × 1080 pixels) and were presented against a black background. Stimuli were presented using PsychoPy2 software (Peirce, 2007; Resource Research Identifier [RRID]: SCR_006571) on an HP desk-top computer running Windows 10.

#### Autobiographical Retrieval Task

Participants retrieved and described autobiographical memories in response to picture cues (**Figure 1**). For each trial, participants were first directed to recall an event corresponding to one of 3 different temporal distances: earlier that same day (Today), 6-18 months ago, or 5-10 years ago. During the initial construction phase of the task (the focus of this manuscript), participants were given a choice of 2 picture cues and were given an 11 second period to select the image they would prefer to use to cue a specific memory. Responses were made via button press with a fiber-optic button box. The screen was replaced with a fixation cross once a response was made, and after the selection period ended, an enlarged version of the selected image was presented in the center of the screen for 5 seconds. Participants used this period to covertly reflect upon a specific event, which was then overtly described in detail throughout the elaboration phase of the task (previously analyzed: Gilmore et al., 2021b, 2021c), consisting of a 116 s response period. Overt responses were recorded with an MRI-compatible microphone. A 2.2 s red fixation cross signaled the end of each trial, and trials were separated by a 19.8 s fixation period. One trial from each of the three time periods was included in each of 6 Autobiographical Recall task scan runs. Prior to scanning, participants were familiarized with the task and practiced retrieving events specific in time and place. For additional details, see Gilmore et al. (2021b).

#### Control Task

An active baseline control task was created to match response demands of the Autobiographical Memory Task without the inclusion of a long-term memory component. Participants were asked to describe the contents of picture stimuli instead of using them to cue specific memories. Trial timings were identical, and participants were first given 11 seconds to select which of 2 picture cues they preferred (control for the construction period of the retrieval task). The screen was replaced with a fixation cross once a response was made, and after the selection period ended, an enlarged version of the selected image was presented in the center of the screen for 5 seconds. Participants used this period to scrutinize the image so that it could be described aloud in detail for the next 116 seconds of description time (control period for the elaboration phase of the retrieval task). A 2.2 s red fixation cross signaled the end of each trial, and trials were separated by a 19.8 s fixation period. Three trials were included per Control Task run, with 2 such runs being collected per participant.

#### fMRI Acquisition Parameters

Data were acquired on a General Electric Discovery MR750 3.0T scanner, using a 32-channel head coil. Functional images were acquired using a BOLD-contrast sensitive multi-echo echo-planar sequence [Array Spatial Sensitivity Encoding Technique (ASSET) acceleration factor = 2, TEs = 12.5, 27.6, and 42.7 ms, TR = 2,200 ms, flip angle = 75°, 64 × 64 matrix, in-plane resolution = 3.2 mm × 3.2 mm]. Whole-brain EPI volumes (MR frames) of 33 interleaved, 3.5 mm-thick oblique slices were obtained every 2.2 s. Slices were manually aligned to the AC-PC axis. A high-resolution T1 structural image was obtained for each subject after functional scans were collected (TE = 3.47 ms, TR = 2.53 s, TI = 900 ms, flip angle = 7°, 172 slices of 1 mm × 1 mm × 1 mm voxels). Foam pillows were provided for all participants to help stabilize head position and scanner noise was attenuated using foam ear plugs and a noise-canceling headset. This headset was also used to communicate with the participant during their time in the scanner. Heart rate was recorded via a sensor placed on the left middle finger and a belt monitored respiration.

### Statistical Analysis

#### fMRI Processing

BOLD timeseries data were processed in AFNI (<u>RRID: SCR_005927</u>). Steps included removal of the first four frames to remove potential T1 equilibration effects (*3dTcat*), despiking (*3dDespike*), slice-time correction (*3dTshift*) and framewise rigid-body realignment to the first volume of each run (*3dvolreg*). Following these initial steps, data from the three echoes acquired for each run were used to remove additional noise using ME-ICA (Kundu et al., 2012) as implemented in the *meica*.*py* AFNI function. This procedure initially calculates a weighted average of the different echo times (“optimally combined” data), which reduced signal dropout and thermal noise and increases contrast-to-noise in each voxel. The resulting image is also registered to the participant’s anatomical image. The individual echo timeseries used to generate the optimally combined data are also submitted to an ICA, and signal decay patterns over time are used to classify components as artifactual in nature (e.g., reflect head motion) or as putatively neural in origin (for more, see Kundu et al., 2012, 2013). Following ME-ICA processing, data were spatially blurred with a Gaussian kernel 3 mm full-width at half-maximum, normalized by the grand mean of each run, and then resampled into 3-mm isotropic voxels and linearly transformed into Talairach atlas space.

#### Audio recording, behavioral response scoring, and alignment of spoken responses to BOLD timeseries data

The protocol for collecting, processing, and scoring spoken response data has been described in detail in prior reports (Gilmore et al., 2021c, 2021b). Briefly, recorded audio was transcribed and scored for content using an adapted form of the Autobiographical Interview (Levine et al., 2002; Gaesser et al., 2011). This procedure separates “Internal” (putatively episodic) details specific to the event details from other types of “External” details. Subcategories of Internal details were expanded from those initially describe by Levine et al. (2002) and included: Activities, Objects, Perceptual, Person, Place, Thought/Emotion, Time, and Miscellaneous. External detail types included Episodic (i.e., details from other events), Repetitions, Semantic statements, and an “Other External” category. Timestamps for each spoken word and phrase were generated and matched with the text in transcripts, and different categories of recalled content were converted into event-related regressors for fMRI data analysis, as will be described below.

#### General Linear Model Analysis

Data were analyzed in AFNI using a general linear model (GLM) approach (*3dDeconvolve*). All task scans consisted of 210 MR frames (214 before initial frame dis-carding) and lasted 7 min, 51 s in duration. Average run-level motion estimates were derived using AFNI’s @1dDiffMag based on three translational and three rotational motion parameters; runs with.0.2 mm/TR were excluded. Within the present data, this resulted in two autobiographical task runs being excluded from four participants and one autobiographical task run excluded from five additional participants. Data were detrended prior to analysis and the analysis was conducted as a mixed block/event related design.

The construction phase (picture selection period) was modeled for each recall condition (today, 6–18 months ago, 5–10 years ago) and the picture description condition using separate regressors with durations of 11 s. The picture display phase for all trials was modeled using a single regressor with a duration of 5 s. Four regressors, each with a duration of 118.2 s, additionally modeled activity associated with the elaboration phase (speaking period) of each autobiographical recall condition (today, 6–18 months ago, 5–10 years ago) and the picture description condition. Effects associated with each category of overtly described internal and external detail during the elaboration phase were modeled across both the autobiographical recall and picture description conditions (i.e., there was a single “place” regressor that accounted for all instances in which a place detail was described during the elaboration period in either of the task conditions) with the spoken duration of each detail included as a duration modulator via AFNI’s dmBLOCK function. Finally, six motion parameters (three translational, three rotational) were included in each subject’s GLM as regressors of non-interest.

#### Hippocampal Region of Interest Analysis

Subject-specific hippocampal masks were generated with Freesurfer (version 6.0; RRID: SCR_001847). Each mask was manually segmented into anterior and posterior long-axis subregions using the uncal apex as a landmark for separation and resampled to the same resolution as the EPI data (example masks can be viewed in **Figure 2A**). Univariate activity was averaged across all voxels in each hippocampal ROI in each subject’s native space for each construction period (picture selection period for Today, 6-18 months, and 5-10 year conditions) relative to the equivalent 11 s picture selection period of the Picture Description control task. A repeated measures ANOVA was conducted on percent signal change relative to the control task, with factors of hemisphere (left, right), long-axis location (anterior, posterior), and temporal distance (Today, 6-18 months ago, 5-10 years ago) as predictors. One-sample t-tests were subsequently conducted to compare the significance of each response versus the Picture Description control task, with both Bonferroni and False Discovery Rate (FDR, Benjamini & Yekutieli, 2001) corrections being reported.

**Figure 2.**
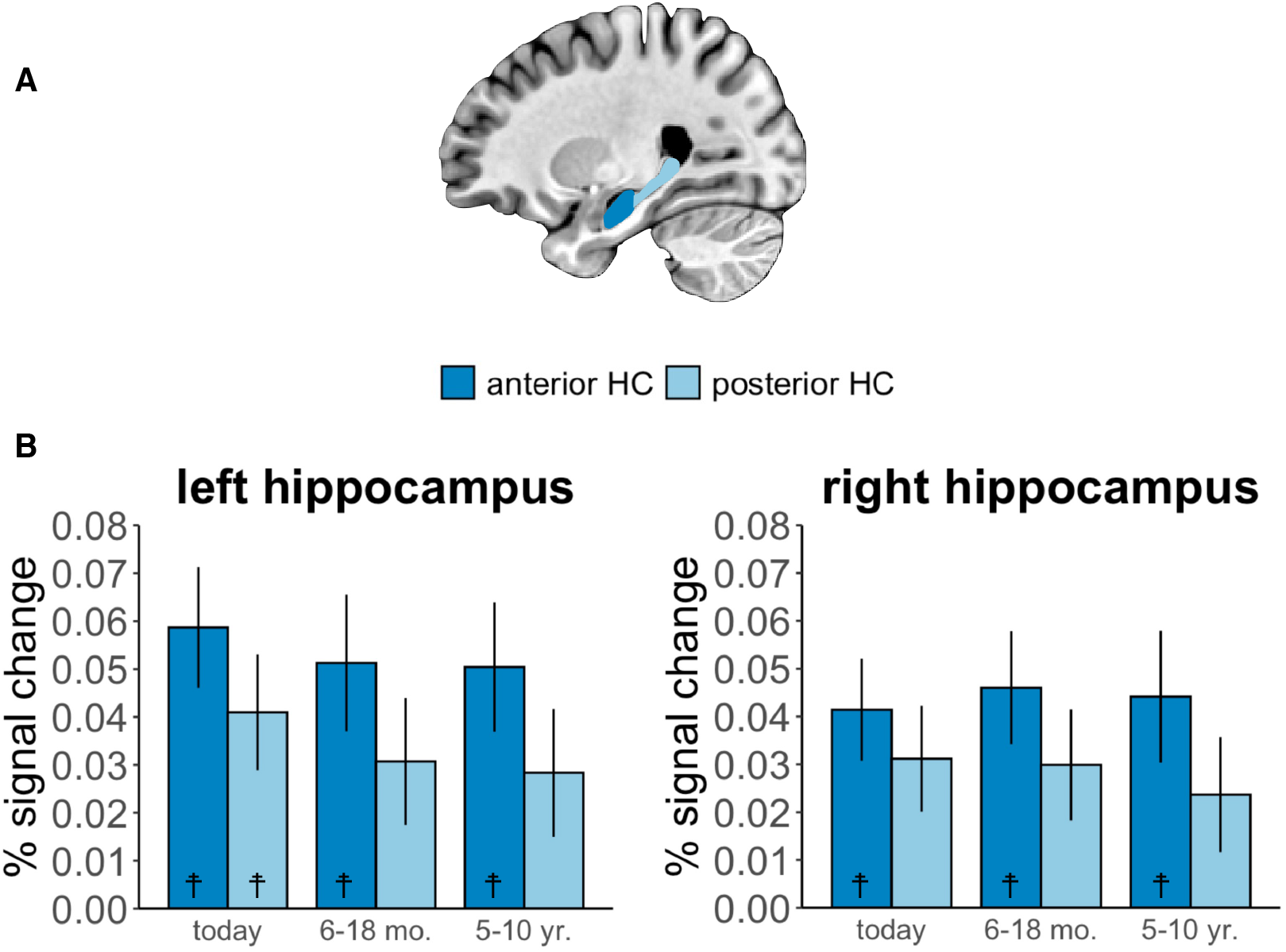
Hippocampal activation during the autobiographical construction phase over time. A) Anterior and posterior hippocampal ROIs, manually segmented for each participant. B) Percent signal change in the anterior and posterior hippocampus during construction of autobiographical memories of varying remoteness relative to the same period of the control picture description task. Error bars reflect standard error of the mean. ☨ p<0.05, construction phase > baseline control task after Bonferroni correction for multiple comparisons. HC=hippocampus, mo.=month, yr.=year.

#### Whole-brain Contrast of Construction and Control Activity

Activity associated with the construction stage was contrasted for autobiographical and control task conditions using a paired-samples, two-tailed *t*-test at the whole brain level. Activity was averaged across all three recall periods for Autobiographical trials. Statistical images achieved a whole-brain *p* < .05 by requiring a voxelwise threshold of *p* < .001 and a minimum cluster extent (*k*) of 15 contiguous voxels as determined using AFNI’s *3dClustSim* and its non-Gaussian (-*acf*) autocorrelation function (Cox et al., 2016). This procedure identified one large cluster spanning multiple lobes as well as the cerebellum. For reporting purposes, discrete regions within this cluster were identified by incrementing the threshold to *t* = 4.55 (p < 5.1 × 10^−5^).

### Code Accessibility

Data analyzed in this report are freely available for download on OpenNeuro.org (Gilmore et al., 2021a).

## Results

A repeated measures ANOVA predicting percent signal change (relative to the control task) as a function of long-axis, hemisphere, and temporal distance revealed a significant main effect of long-axis (F(1,39)=7.76, p=0.008, η^2^=0.013), which was driven by greater activation in the anterior than posterior hippocampus (**Figure 2B**). There was no significant effect of temporal distance (F(2,78)=0.14, p=0.87, η^2^=0.001) and no interaction between long-axis and temporal distance (F(2,78)=0.70, p=0.50, η^2^=0.0004). There was also no main effect of hemisphere (F(1,39)=1.64, p=0.21, η^2^=0.002) or interaction between hemisphere and the other variables in the model (all p>0.097). Thus, during the memory construction stage, the anterior hippocampus was more active than the posterior portion, and there was no evidence of a temporal gradient over time.

Construction-related hippocampal activity was then compared to the baseline task using one-sample t-tests to identify which conditions, if any, resulted in significant hippocampal activation. Following Bonferroni correction for multiple comparisons (requiring *p* < 0.004), the anterior hippocampus was significantly activated for all temporal distances, in both the right (today: t(39)=3.88, p=0.0004; 6-18 months: t(39)=3.89, p=0.0004; 5-10 years: t(39)=3.21, p=0.0027) and left (today: t(39)=4.67, p=0.00004; 6-18 months: t(39)=3.59, p=0.0009; 5-10 years: t(39)=3.73, p=0.0006) hemispheres. The only significant posterior hippocampal effect was in the left hemisphere for the “today” condition, t(39) = 3.39, p = .0016 (other p’s > .008). Applying a more liberal False Discovery Rate correction for multiple comparisons (Benjamini & Yekutieli, 2001) only resulted in a single change, with right posterior hippocampal activity for “today” now also significant. Thus, during memory construction, we observed robust activation in the anterior hippocampus regardless of memory remoteness.

Unlike the later elaboration phase, no overt behavior outside of a button press was available during the construction phase. Therefore, to contextualize the hippocampal findings, a whole-brain voxelwise contrast was conducted to compare neural activity during the construction phase of autobiographical retrieval with that of the matched control task, averaging across temporal distances (requiring voxel-wise p<0.001, k≤15, to achieve a whole-brain p<.05). In addition to observing significant activity in anterior hippocampal regions, we observed strong activation in regions typically associated with autobiographical recall, including medial prefrontal, posterior cingulate, lateral temporal, and parahippocampal cortices as well as the angular gyrus (**Figure 3; Table 1**; (Svoboda et al., 2006; Spreng et al., 2009), collectively supporting a retrieval-based interpretation of the present hippocampal findings.

**Table 1.** Peak regions identified in the voxel-wise analysis of construction phase > control task activity. Coordinates refer to centers of mass in MNI space. Primary clusters were identified at *t* > 3.55; Discrete peaks identified in the posterior parietal-occipito-temporal-cerebellar cluster were separated at a *t* > 4.55.

| Region | X | Y | Z | Cluster Size | Peak T |
| --- | --- | --- | --- | --- | --- |
| **Construction phase > control task** |  |  |  |  |  |
| Posterior parietal-occipito-temporal-cerebellar cortex | 2 | -65 | -1 | 2806 | 10.02 |
| Bilateral posterior cingulate/ventral medial parietal cortex | -1 | -54 | 15 | 456 | 10.02 |
| Right occipital/fusiform cortex | 34 | -81 | -4 | 311 | 7.09 |
| Left visual cortex | -24 | -87 | -7 | 264 | 6.95 |
| Left parahippocampal cortex | -24 | -40 | -17 | 56 | 6.55 |
| Right parahippocampal cortex | 28 | -38 | -17 | 38 | 6.91 |
| Right Cerebellum (Crus 1) | 8 | -79 | -21 | 24 | 5.49 |
| Left anterior hippocampus | -20 | -17 | -18 | 19 | 5.91 |
| Left posterior fusiform | -40 | -69 | -18 | 17 | 5.04 |
| Left middle frontal gyrus/superior frontal sulcus | -24 | 24 | 51 | 240 | 6.31 |
| Bilateral medial prefrontal cortex | -1 | 56 | -13 | 199 | 7.30 |
| Left angular gyrus | -43 | -72 | 34 | 173 | 6.93 |
| Left anterior frontal cortex | -17 | 63 | 10 | 109 | 6.65 |
| Right cerebellum (lobule IX) | 8 | -52 | -39 | 78 | 6.08 |
| Right middle temporal gyrus | 59 | -6 | -19 | 78 | 6.07 |
| Left middle temporal gyrus | -56 | -3 | -16 | 60 | 6.15 |
| Left inferior frontal sulcus | -43 | 19 | 27 | 59 | 5.19 |
| Left supplementary motor area | -1 | 18 | 48 | 44 | 5.03 |
| Right angular gyrus | 43 | -69 | 33 | 38 | 5.33 |
| Right anterior hippocampus | 24 | -18 | -21 | 26 | 5.72 |
| Brainstem | -1 | -28 | -17 | 24 | 5.62 |
| Left inferior frontal gyrus (pars orbitalis) | -36 | 38 | -12 | 24 | 5.46 |
| Left superior occipital gyrus/cuneus | -14 | -95 | 33 | 15 | 4.18 |
| **Control task > Construction phase** |  |  |  |  |  |
| Right anterior intraparietal sulcus | 43 | -43 | 45 | 52 | -5.07 |
| Left supramarginal gyrus | -59 | -40 | 42 | 30 | -5.51 |
| Right inferior frontal gyrus (pars triangularis) | 50 | 42 | 2 | 15 | -5.39 |

**Figure 3.**
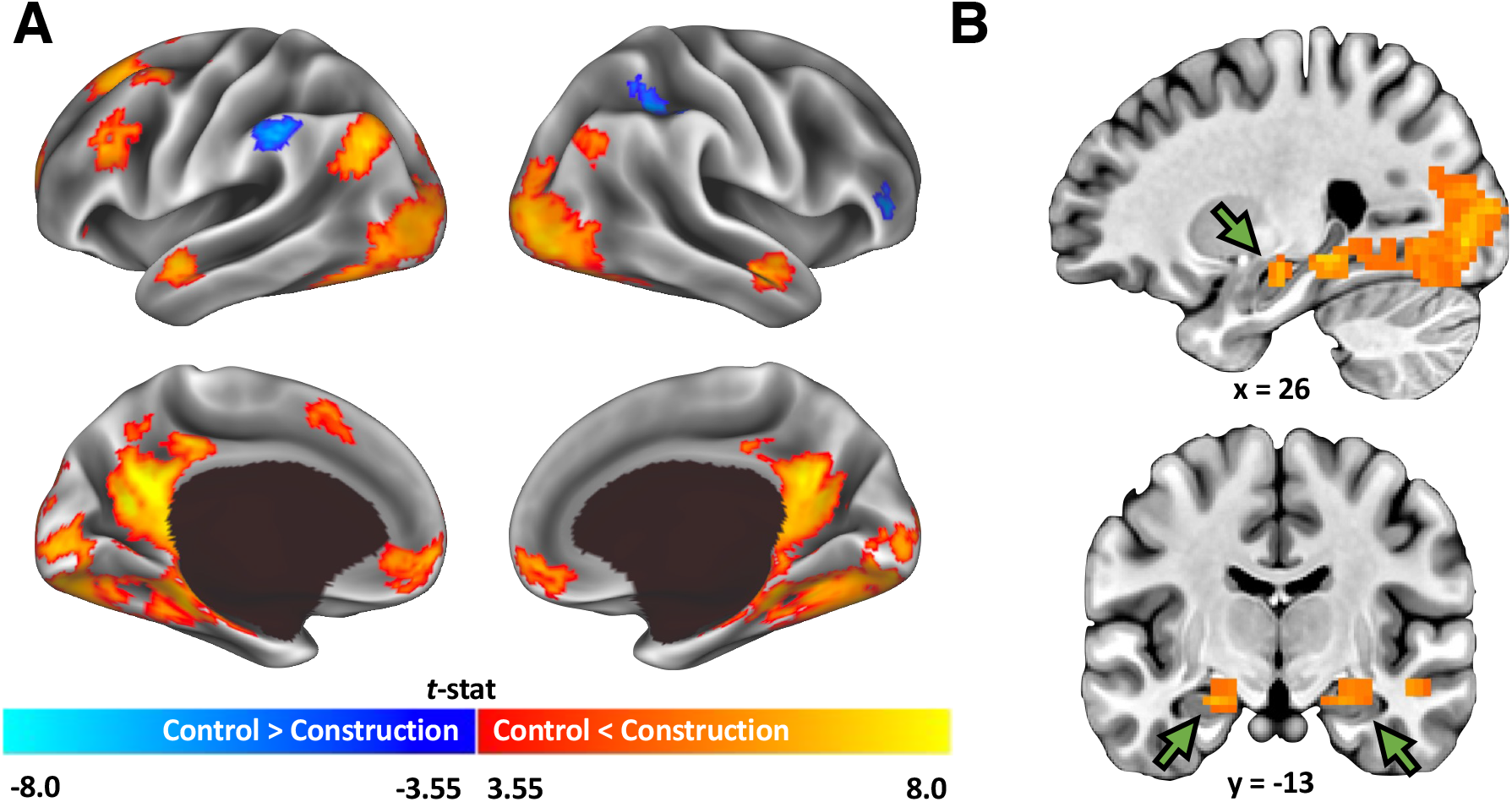
Whole-brain univariate analysis of the memory construction phase. A) Surface reconstruction of cortical regions with significant activation during memory construction relative to the same period of the control task, averaged across temporal distances. Data were surface-projected using Connectome Workbench (software (Marcus et al., 2011). B) Sagittal and coronal slices of the same contrast displaying activations identified in the anterior hippocampus, as pointed to by the green arrows.

## Discussion

This work aimed to understand hippocampal contributions to autobiographical memory retrieval during an initial construction stage by taking both temporal distance and long-axis location into account. We found that the anterior hippocampus is active during the construction period of autobiographical memory retrieval, regardless of memory age. Notably, neither the standard model of consolidation nor trace transformation theory emphasizes the component processes of autobiographical retrieval (i.e., construction vs. elaboration) as determining features of hippocampal involvement in retrieving consolidated memories, although the current results suggest that these, along with long-axis placement, are critical aspects in deriving an answer.

A key question that arises from, but is not specific to the current results is how one can unpack what is occurring during the construction stage given the lack of overt behavior. Indeed, prior work with these data focused on elaboration specifically because overt speech could be leveraged to understand dynamic processes related to retrieval (Gilmore et al., 2021c, 2021b). Presumably, participants in this study used the picture selection period to extract meaning from the picture cue, and then search for and retrieve a memory for a related event that they could proceed to elaborate upon, but we did not ask participants what they were doing during the construction period. Indeed, it is unclear how to probe the subjective experience of construction at the trial-level without changing participant behavior. However, our whole-brain univariate analysis of recall during the construction stage relative to the control picture description task revealed canonical autobiographical memory regions, supporting the interpretation that the picture cues in the construction period elicited autobiographical retrieval in preparation for elaboration. An alternative account might be that, rather than a construction-like process as predicted by Conway (2005), the activity instead represents direct retrieval of a memory as a function of cue/trace overlap, in line with the encoding specificity principle (Tulving and Thomson, 1973).

We do not yet understand the mechanisms, common or distinct, that support retrieval as it unfolds across multiple trial stages. In general terms, it is possible that during memory construction the anterior hippocampus retrieves the central aspects of memory, which is used as a scaffold for details retrieved during elaboration. This idea fits with Conway’s Self-Memory System and its proposal that autobiographical memory is organized hierarchically, such that event-specific details are contextualized by general event structures (Conway, 2009; Conway & Pleydell-Pearce, 2000; Sheldon et al., 2019). There is evidence that when a specific autobiographical memory is elicited with a thematic cue (e.g. “basketball game”), the first thoughts tend to be that of general autobiographical knowledge pertaining to the event followed by event-specific details (Haque and Conway, 2001). This observation suggests that general aspects of autobiographical events (perhaps retrieved by the anterior hippocampus in concert with frontal regions) may be used to access more specific ones (perhaps retrieved by the posterior hippocampus and posterior cortical regions) during intentional retrieval (Conway and Pleydell-Pearce, 2000; Conway, 2009). The question of whether this process must proceed from general to specific in the hippocampus, or rather if it can occur in the opposite direction (Maurer & Nadel, 2021) or in parallel (Gilboa and Moscovitch, 2021), remains to be fully addressed. The way in which retrieval is cued is likely important. For example, there is evidence that orienting participants to conceptual or contextual information during retrieval recruits anterior and posterior memory systems respectively, providing some support for parallel processing depending on memory content (Gurguryan and Sheldon, 2019; Sheldon et al., 2019).

Reconciling the current observation of significant anterior hippocampal activity for both recent and remote events during construction, with prior observations of temporally graded posterior hippocampal activity during elaboration/overt recall, is important. Especially so, when one considers the types of information that the anterior and posterior hippocampus are proposed to represent, namely, gist and detail respectively (Poppenk et al., 2013; Sekeres et al., 2018; Audrain and McAndrews, 2020). As described in our previous work, even remote memories were retrieved with rich detail in our sample (Gilmore et al., 2021b). One possibility is that the anterior hippocampus represents more detail than is typically ascribed to it, or else comes to support retrieval of such details over time as they become integrated and bound to central aspects of the memory with consolidation. Perhaps what is retrieved by the anterior hippocampus during the construction period is not a scaffold devoid of specific detail but rather the representative details of the memory such as who, what, where, and when. Indeed, the anterior hippocampus plays an active role in constructing scenes (Zeidman and Maguire, 2016), which some might consider more “detailed” than “gist-like”, and it has also been associated with the recombination of specific event details in the service of simulating future events (Addis et al., 2011; Addis and Schacter, 2012). The posterior hippocampus may be additionally important for details retrieved when one “replays” memory or projects oneself through memory in a more continuous “frame-by-frame” spatiotemporal fashion, which is a hallmark of mental time travel (Tulving, 1985; Suddendorf and Corballis, 2007), and which would necessitate fine-grained shifts in representational content (Takehara-Nishiuchi, 2020). In this vein, emphasizing retrieval of perceptual or peripheral detail as a uniquely posterior hippocampal function may be incorrect, and could account for discrepancies reported in the literature ascribing detail representation to the anterior hippocampus (Zeidman and Maguire, 2016; Tompary and Davachi, 2017; Dandolo and Schwabe, 2018; Cowan et al., 2021). It follows that defining what constitutes “gist” and “detail” is critical for precisely delineating how anterior and posterior hippocampal segments differently support memory along these dimensions. Indeed, while subjective vividness has historically been used as an index of detail retrieved in covert memory paradigms, recent evidence suggests that vividness may track more closely with the gist of an event (defined as memory for names of people, objects, and places comprising the event) than nuanced perceptual detail (Cooper and Ritchey, 2022), highlighting that a consensus regarding this terminology is paramount.

To conclude, findings from this dataset—both the current results and those reported previously—can be taken to support either of two dominant views of hippocampal contributions to remote retrieval depending on analysis choices (task- and neuroanatomy-related components). Given the decades-long debate centering on the role of the hippocampus in recent and remote memories, perhaps this is appropriate. Moving forward, it will be critical to articulate the sub-processes involved in construction, as well as to come to a consensus regarding what constitutes gist and detail. Precise delineation of these concepts will inform testable predictions regarding the role of the anterior hippocampus in construction over time. The data presented here highlight that the stage of retrieval and anatomical location along the long-axis of the hippocampus analyzed are critical considerations not fully integrated into current models. Our understanding of anterior and posterior hippocampal contributions to retrieval of autobiographical memory over time can only be enhanced by understanding the component processes inherent to such retrieval.

## Acknowledgements

This work was supported by the National Institute of Mental Health (NIMH) Intramural Research Program (ZIA MH002920) and by NIMH Grant R01 MH060941 (to DLS). Thank you to Sarah Kalinowski, Alina Quach, and Bess Bloomer for assistance with collection and preparation of the data.

